# Loss of MGST1 during fibroblast differentiation enhances vulnerability to oxidative stress in human heart failure

**DOI:** 10.1101/2025.11.14.688417

**Authors:** Mohamad Youness, Onne A.H.O. Ronda, Ankit Pradhan, Filip Rega, Karin R. Sipido, H. Llewelyn Roderick

## Abstract

**Background:** Oxidative stress (OS) resulting from an imbalance between reactive oxygen species (ROS) production and antioxidant defences is a major mechanism exacerbating HF, involving activation of cardiac fibroblasts (FB) and fibrosis. In addition to reducing ROS production and/or scavenging, boosting cellular anti-oxidant capacity is a potential strategy, with FB an interesting target.

**Methods:** We interrogated single-nucleus RNA sequencing (snRNA-seq) data from human failing and non-failing hearts to identify OS modulating genes. Differential gene expression analysis in FB and comparison with other cell types was used for identification of candidate genes. The functional role of the identified candidate was studied in primary isolated human FB in culture, using knock-down strategies and phenotype manipulation using TGF-β1.

**Results:** Across the different cardiac cell types, FB showed the greatest enrichment for OS-modulating genes downregulated in HF. Of these genes, the microsomal glutathione S-transferase 1 (MGST1), was exclusively expressed and downregulated in FB, inversely correlating with elevated ROS levels in HF tissue. TGF-β1 treatment of non-HF FB reduced MGST1 expression. MGST1 knockdown in non-HF FB raised expression of periostin, but not of collagen. ROS production and susceptibility to oxidative damage and lipid peroxidation were also increased. Downregulation of ferroptosis-suppressing and iron-handling genes inversely correlated with MGST1 expression.

**Conclusion:** Loss of MGST1 in HF is unique to FB, linked to FB activation. It impairs FB antioxidant capacity, exacerbating oxidative stress, and reduces resistance to ferroptosis. Through these mechanisms, loss of FB MGST1 induces a deleterious positive feedback on cardiac remodelling in human HF.

## Introduction

Over the past decades, clinical and experimental studies have generated substantial evidence indicating that oxidative stress, i.e. overproduction of reactive oxygen species (ROS) (ROS species here) and/or dysfunction of the antioxidant defence systems, contributes to adverse cardiac remodelling involving cardiomyocyte hypertrophy and death and fibrosis and its subsequent progression to HF (1–4). While in physiology ROS are key signalling mediators, in pathology, redox imbalance leads to peroxidation of proteins and lipids, DNA damage, and misregulation of signalling that together result in cellular dysfunction, irreversible damage and death. While CM are established as a major source of ROS in the heart emanating from compromised mitochondria for e.g. in HF, these short-range bioactive molecules are generated and function in all cell types of the heart, suggesting their therapeutic targeting. However, effective therapies are yet to be identified.

In cardiomyocytes, excessive ROS can directly impair contractile function by modifying proteins essential for excitation-contraction coupling, as well as triggering hypertrophy and inducing gene expression. Oxidative stress (OS) is also central to endothelial dysfunction in HF contributing to atherosclerosis development. ROS signalling is further strongly linked with cardiac fibrosis where it is increased downstream of angiotensin signalling. Angiotensin is a potent activator of FB, that stimulates transition of FB from their quiescent basal state to a contractile myoFB state and induces transcription and translation of extracellular matrix (ECM) components contributing to fibrosis (reviewed in (5)).

Feed-forward mechanisms between TGF-β and ROS are well documented, where ROS increases release of TGF-β into its active form, while the latter enhances mitochondrial ROS production and decreases anti-oxidant defences (6). Additionally, ROS further modulates matrix degradation through matrix metalloproteinases (MMPs) expression (7,8). ROS brings about these changes in expression of ECM components by modulating the activity of several transcription factors -most notably through the indirect activation of HIF1α, which enhances the transcription of hypoxia-responsive genes (9).

OS and inflammation are interrelated and mutually re-enforcing (6,10,11). The activation of cardiac fibroblasts (FB) in response to inflammatory, neurohormonal, or mechanical cues, leads to their migration, proliferation and transition into new states driving cardiac fibrosis (12–14). Interestingly, beyond their role in maintaining matrix and providing structural integrity of the heart, FB have been recently shown to protect cardiomyocytes during myocardial infarction from ferroptosis (a non-apoptotic and iron-dependent form of regulated cell death) through paracrine effects and direct cell-cell interaction (15). Hence, FB protection from OS is beneficial in reducing cardiac fibrosis and in protecting cardiomyocytes from death.

Although preclinical studies found that reducing ROS and/or augmenting the heart’s own antioxidant defences improved outcome after cardiac injury and progression to HF (16–18), clinical translation of these findings was mostly disappointing, with possibly some benefit of increasing anti-oxidant capacity (reviewed in (4)). New avenues for improving anti-oxidant therapies include a more targeted approach to specific cell types and molecular pathways through nanomedicine, as pioneered in other organs (19–21).

To investigate the role of ROS imbalance in FB in human HF, we leveraged our recent snRNA-seq dataset from human failing (HF) and non-failing (NF) hearts, and probed for FB-specific alterations in expression of genes involved in the OS response and for candidate targets involved in anti-oxidant defence. Relative to the other major cell types of the heart, we find FB to show the greatest alteration in genes involved in anti-oxidant responses. Moreover, we identify the microsomal glutathione S-transferase 1 (MGST1) as exclusively expressed in FB in myocardial tissue, with a decrease in its expression in HF regardless of aetiology. We explore the relationship between MGST1 and inflammation by investigating whether TGF-β pathway activation of FB regulates its expression. Finally, we show through intervention studies that loss of MGST1 increases vulnerability of primary human cardiac FB to OS.

## Methods

An extensive methods section is provided as Supplementary Methods. We used previously generated snRNA-seq data from NF (n = 4) and HF hearts (ischemic cardiomyopathy [ICM], n = 5; dilated cardiomyopathy [DCM], n = 6) reported by Youness et al (in press) (22), as well as tissue samples from the same hearts. Fibroblasts for functional studies were isolated from NF hearts. Imaging assays included dihydroethidium (DHE) staining in tissue, CellROX-Green and ThiolTracker for cellular ROS and GSH, and the Image-iT Lipid Peroxidation sensor (quantified in CellProfiler); MGST1 knockdown and immunostaining/immunoblotting protocols are described in the Supplementary Methods. A publicly available bulk RNA-seq of primary cardiac FB ± TGF-β1 (GSE225336) was reanalysed for MGST1 expression (23). For statistical testing, cells isolated from each heart and cultured in the same flask prior to distributing between experimental conditions are considered a separate biological replicate (N), with each condition considered paired/repeated measurements. Statistical tests used in each analysis are reported in the figure legends. GraphPad Prism software (10.2.3) was used for statistical testing.

## Results

### FB exhibit the greatest downregulation of anti-oxidant genes in HF

Within our snRNA-seq dataset generated on 16 HF tissue samples and 4 NF samples, we explored which cell type exhibited the greatest remodelling in expression of genes responsive to ROS. To this end, we scored individual nuclei of the major cardiac cell types for their expression of a curated list of ROS-responsive genes (*HALLMARK_REACTIVE_OXYGEN_SPECIES_PATHWAY* gene set from the Molecular Signatures Database (MSigDB) Hallmark collection) (other cell types in Suppl.Fig.1a). In NF, FB had the highest module score relative to other cell types. This score was however significantly decreased in HF, highlighting the loss of expression of genes involved in the ROS response.

Having identified FB as the cell type that exhibited the greatest remodelling of this curated list of ROS responsive genes, we next set out to identify ROS-responsive genes that were downregulated in HF FB. To this end, we performed a pseudobulk differential gene expression analysis on NF and HF FB. This analysis identified MGST1 and GPX3, encoding proteins known for their anti-oxidant activity, among the top downregulated genes in HF (Fig.1b). A heatmap representation of these differentially expressed ROS responsive genes between HF and NF in FB illustrates the loss of anti-oxidant protection capacity in HF FB (Fig.1c). The most highly expressed genes in FB are MGST1, PRDX6 and GPX3, and of these three, MGST1 is the only one that is FB-specific.

**Fig. 1.**
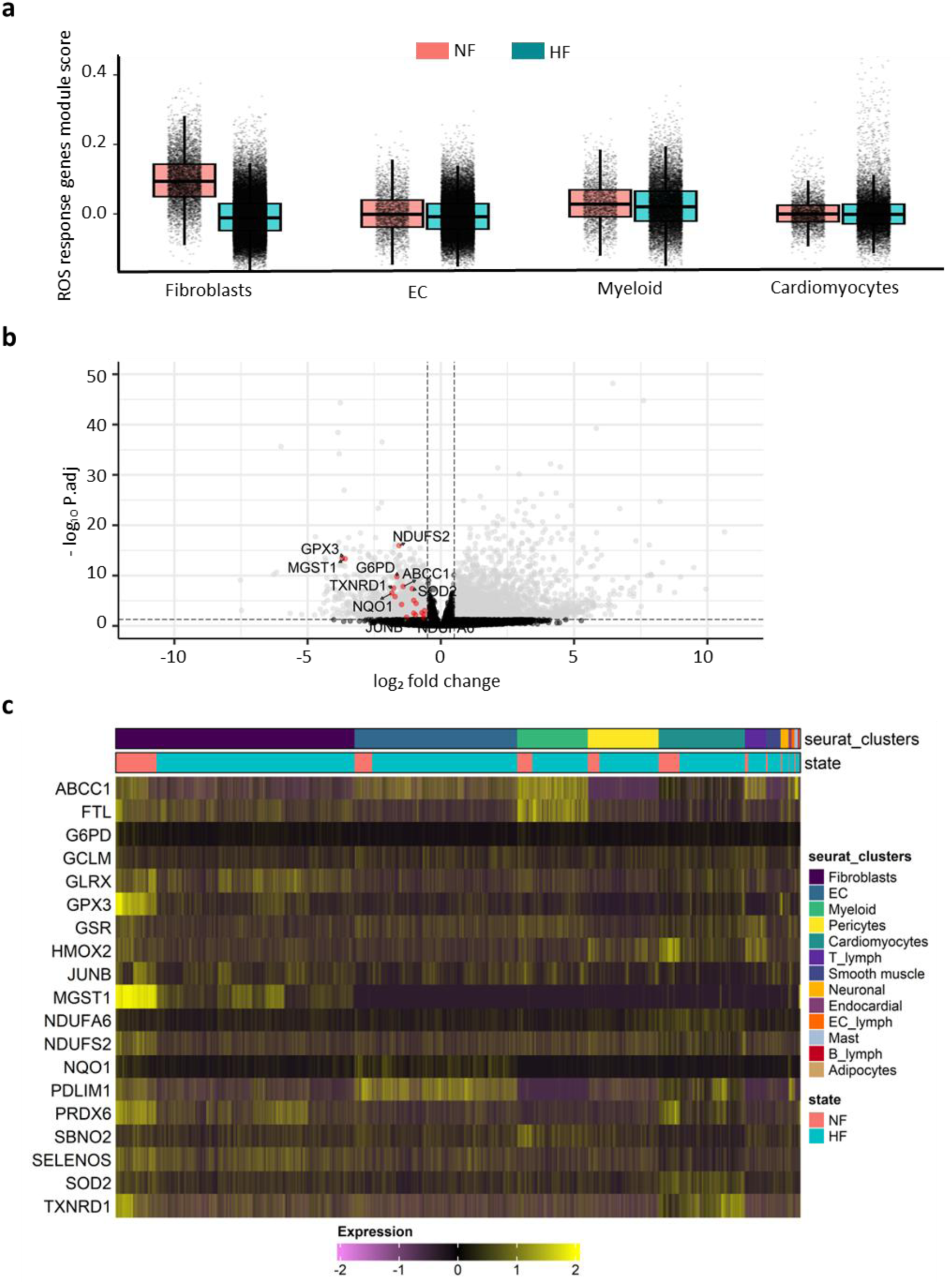
Loss of anti-oxidant capabilities in HF predominates in FB. **a**, Scoring of ROS response gene expression in main cardiac cell types. Genes taken from HALLMARK_REACTIVE_OXYGEN_SPECIES_PATHWAY (M5938). **b,** Volcano plot of gene expression changes in HF FB versus NF. Grey dots represent all genes that are differentially expressed with absolute log2 fold change (> 0. 5) and adjusted p-value thresholds (0.05). Red dots represent ROS-responsive genes (from Fig.1a) that are downregulated in HF. Differential gene expression was done using DESeq2 (pseudo-bulk). **c**, Heatmap showing the scaled gene expression of the ROS-responsive genes scored in **a** that are also downregulated in HF.

### MGST1 downregulation and tissue ROS are anti-correlated in the failing human heart

We next studied the expression and function of MGST1 in NF and HF cardiac tissue. Supporting the snRNAseq findings, and taking advantage of the FB specific nature of its expression, RT-qPCR analysis of RNA prepared from an extended collection of human midmyocardial tissue samples identified downregulation of MGST1 in HF (Fig.2a), irrespective of disease aetiology (Suppl.Fig.2a).

**Fig. 2.**
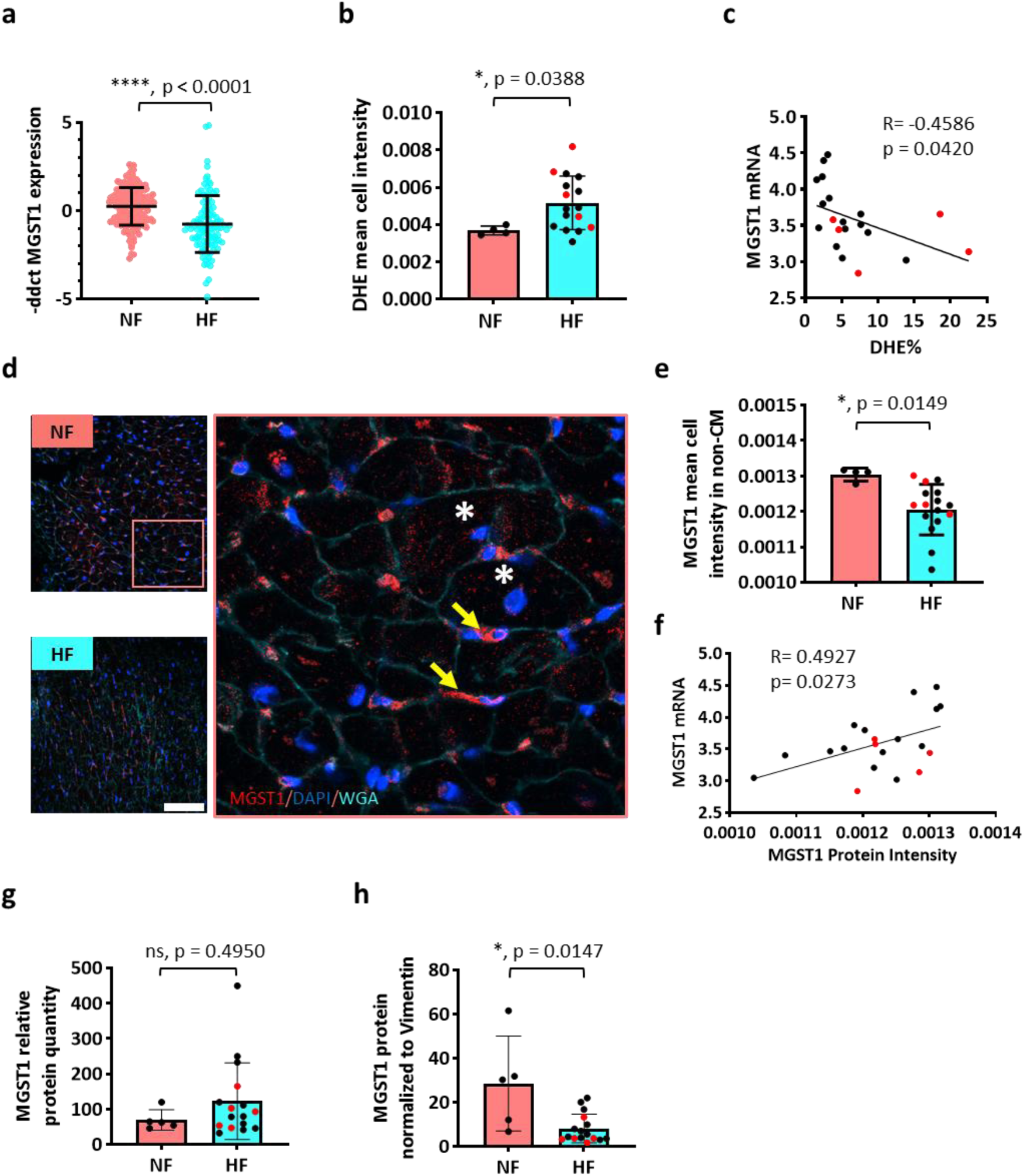
Loss of MGST1 in the failing heart. **a.** Relative mRNA expression of MGST1 in human cardiac tissue demonstrating decrease in MGST1 expression in HF (N= 104) compared to NF (N=166). Unpaired t-test. **b.** Detection of superoxide in myocardium by DHE staining. Compared with NF (N=4), the mean cell fluorescence intensity of DHE in HF (N=16) was significantly increased. Mann-Whitney test was used. Error bars represent mean ± SD. **c.** Pearson correlation of MGST1 mRNA level with % DHE from fig.2b (matched samples). **d.** MGST1 fluorescence staining (red) of NF and HF myocardial tissue sections. Nuclei are stained with Hoechst 33258 (blue) and membranes stained with WGA (cyan). Right is a zoomed image highlighting the positive staining of MGST1 in interstitial cells (non-CM; arrows) and not in CM (* in the centre of CM). Scale bar = 100 µm. **e.** MGST1 fluorescence staining quantification of non-cardiomyocytes (see methods). Unpaired t-test. **f.** Pearson correlation of MGST1 mRNA level and MGST1 protein level intensity (matched samples). **g.** Total MGST1 protein abundance in NF (N=5) and HF (N=16) cardiac tissue lysates determined by Mass Spectrometry (same samples used for snRNA-seq and histology). Mann Whitney test. **h.** Normalized MGST1 protein abundance in NF and HF cardiac tissue lysates (same samples used for snRNA-seq and histology) relative to the mesenchymal marker Vimentin. Mann Whitney test. Error bars represent mean ± SD. ns, no significance. Red dots represent ICM-scar samples.

To validate MGST1 expression at protein level, we performed immunofluorescence labelling in the same samples that were used for snRNA-seq data (Fig.2d). MGST1 showed a pattern of expression in the interstitial cells (arrows) that was visually lower in intensity in HF. To further quantify this expression in these interstitial cells, we measured mean fluorescent intensity in these areas, excluding cardiomyocytes (marked by asterisk, see Suppl.Fig.2f & Suppl.Methods for image processing). This analysis confirmed that MGST1 was downregulated in HF tissue sections (Fig.2e). The MGST1 protein expression was positively correlated with MGST1 mRNA expression from the same samples, further validating our results (Fig.2f).

We also analysed tissue MGST1 expression by immunoblotting but expression could not be detected (data not shown) possibly due to the low mass/cell volume of FB (relative to cardiomyocytes) combined with low intracellular expression. Consequently, we performed mass spectrometry analysis on tissue lysates and could confirm the presence of MGST1 protein, although standard quantitation relative to total peptides detected did not show a difference between HF and non-HF tissues (Fig.2g). Given that MGST1 is expressed in FB, we next normalised its expression to vimentin, a marker of mesenchymal cells, and found that MGST1 levels were significantly downregulated in HF compared to non-HF samples, suggesting a reduction within the mesenchymal cell population (Fig.2f).

Given that decreased MGST1 should be associated with increased ROS, using DHE as an indicator of superoxide, we next explored in matched samples to those used for snRNA-Seq, whether tissue-level ROS was increased (Suppl.Fig2b). Overall DHE mean cell intensity was higher in HF tissues (Fig.2b), across aetiologies. Similar findings were obtained when total area of DHE staining was quantified (Suppl.Fig.2c-e). The intensity of DHE staining negatively correlated with MGST1 mRNA levels, supporting the hypothesis that MGST1 contributes to anti-oxidant activity in HF (Fig.2c).

### MGST1 expression is downregulated in activated HF FB

In the evolution of HF, FB transition from resting (aka basal or quiescent) states through several intermediate states to one or more activated states. These states and their differences between NF and HF can be identified as distinct clusters in the snRNAseq dataset (12,22,24–26) (Fig. 3b). Our previous analysis identified states FB1, 4 and 5 as representing activated states enriched in HF and FB0, 2 and 3 being as dominant in NF, representing resting FB states. Notably, and supporting its elevated expression in NF, FB states 0, 2 and 3 are most enriched for MGST1. Accounting for the decreased MGST1 in HF, enriched states (activated FB1, 4 and 5), do not express significant MGST1 and basal sates (FB 0, 2 and 3) show diminished expression (Fig. 3b, c; Suppl.Fig.3a). Since states 1, 4 and 5 arise from the basal states 0, 2 and 3, these data support the notion that loss of MGST1 correlates with FB activation. This hypothesis is further supported by the anti-correlation of the pseudo-time trajectories of MGST1 with FB activation markers POSTN, ACTA2, COL1A1 and FAP (Fig.3c).

**Fig. 3.**
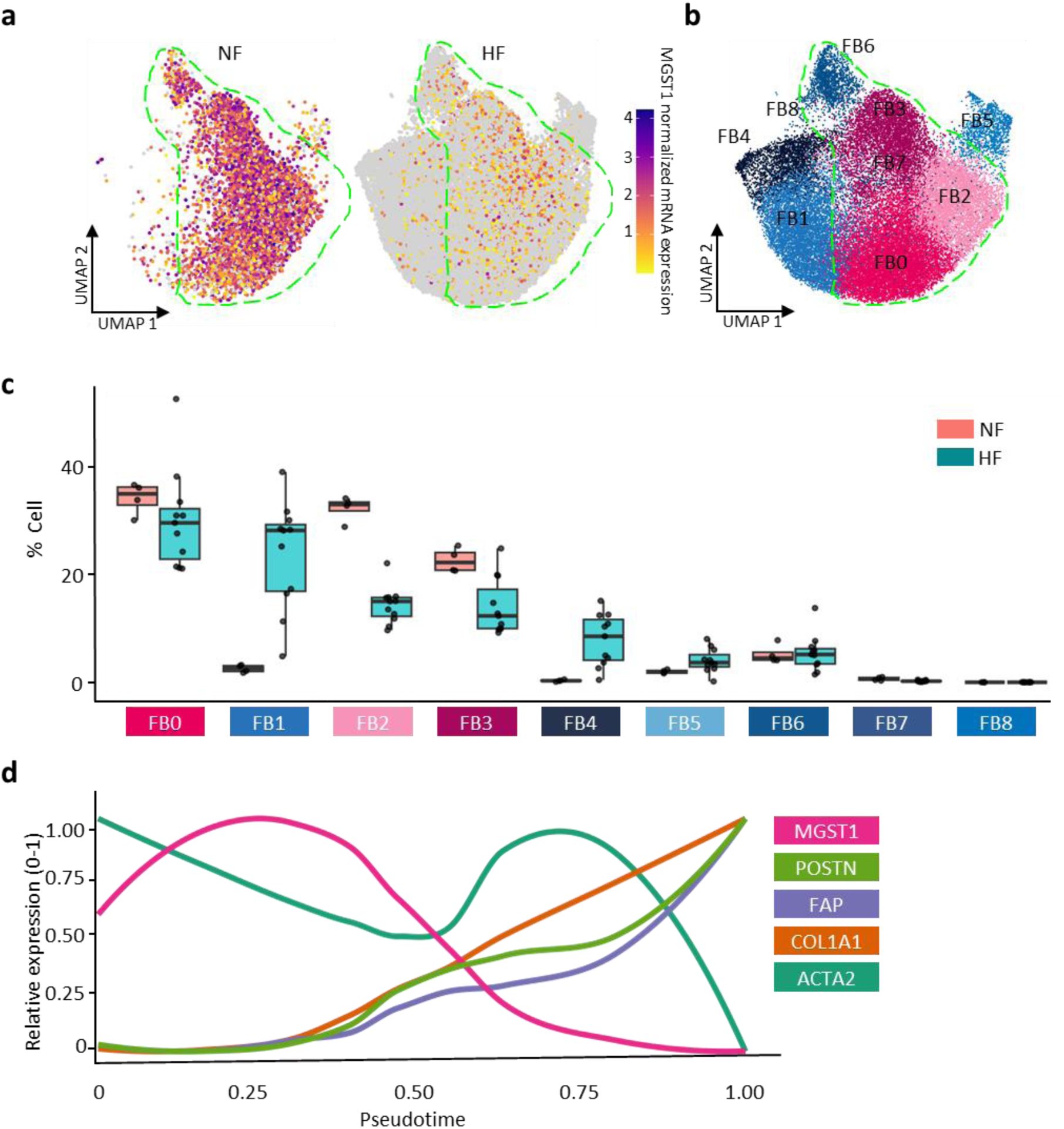
MGST1 is downregulated in activated FB. **a.** Normalized expression of MGST1 in NF and HF FB. Data represents FB clusters taken from snRNA sequencing dataset (Youness et al). Dotted green line represents the MGST1+ cells in NF. **b.** 2D visualisation (UMAP) of FB subclusters, highlighting different FB states. The dotted green line represents the MGST1+ cells in NF representing clusters FB0,2,3, and 6. **c.** Relative abundance of each FB cluster in HF and NF, as measured in the integrated snRNA-seq dataset. **d.** Relative expression levels of selected genes along pseudotime (MGST1, COL1A1, FAP, POSTN, and ACTA2). Normalized expression is bounded by min-max (0-1) to show the trend of expression.

### Activation of human cardiac FB by TGF-β1 leads to a reduction in MGST1

To explore a potentially causal relation between FB activation and MGST1 expression, we examined these features in NF human cardiac FB exposed to TGF-β, a key driver of fibroblast activation and differentiation. Data derived from a publicly available RNA-seq dataset of the time course response of human cardiac FB to TGF-β1, indicated downregulation of MGST1 mRNA by 24 h post-treatment, which remained suppressed for the three-day duration of the experiment (Suppl.Fig.4a).

We then assessed this link further in primary cultures of FB isolated from NF hearts exposed for 4 days to TGF-β1 alone or in combination with SD-208, a selective TGF-β receptor I (TGF-β-RI) kinase inhibitor. NF FB similarly cultured for 4 days but without treatments (only DMSO vehicle) were used as control. Confirming the activation of FB by TGF-β1 and its inhibition by SD-208, mRNA markers of FB activation were significantly induced by TGF-β1 and inhibited by SD208. Consistent with a basal activation of TGF-β-RI kinase in FB cultures, co-application of SD208 reduced expression of activation markers below that observed in controls (Fig.4a).

**Fig. 4.**
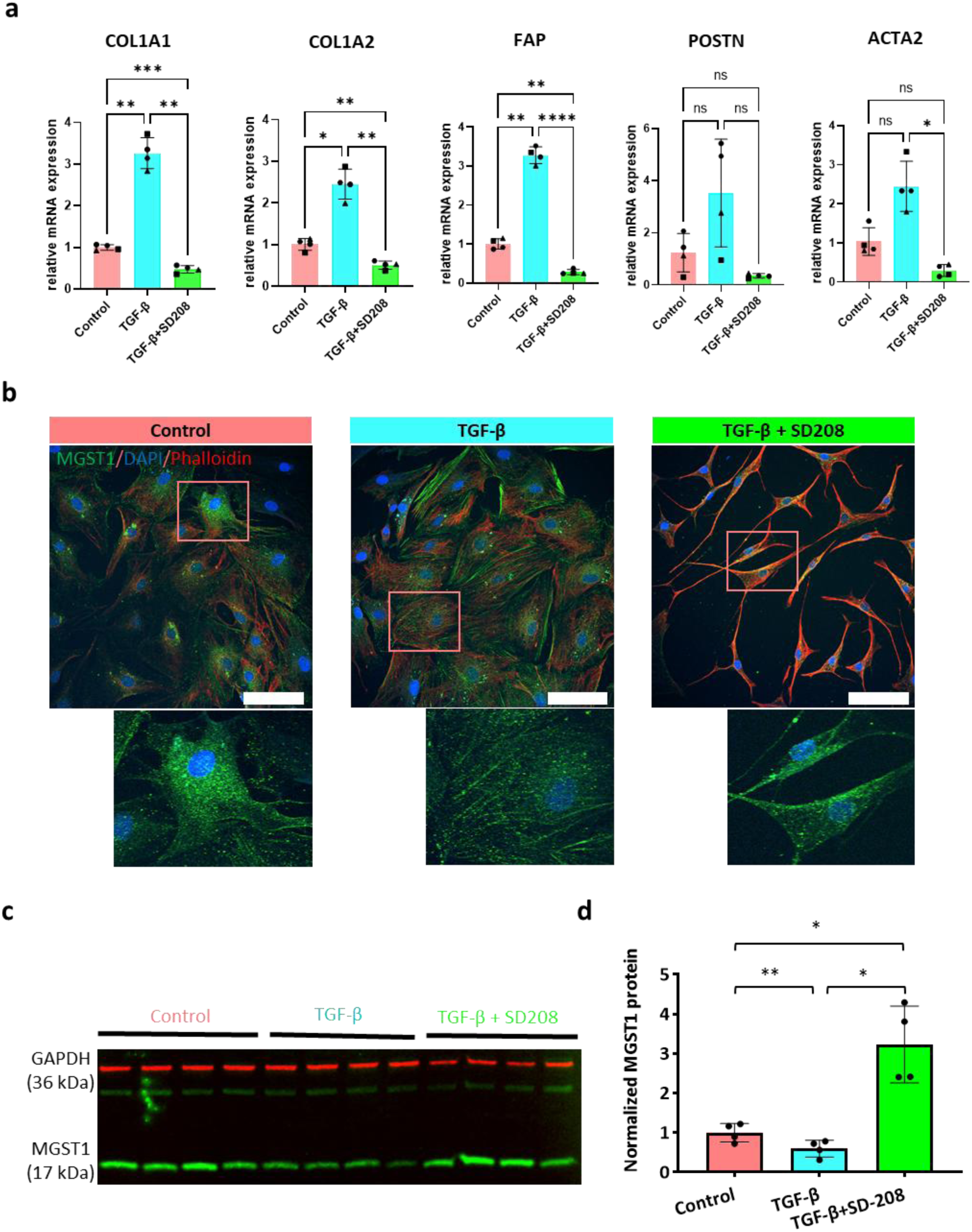
Activation of human cardiac fibroblasts by TGF-β leads to a reduction in MGST1. **a.** Relative mRNA expression level of fibrosis and FB activation-related genes upon MGST1 silencing. One-way ANOVA (RM) was used with Tukey’s correction for multiple comparisons. **b.** Representative image of MGST1 fluorescent staining (green) of NF donor FB cells without TGF-β (left), with TGF-β (middle) or with TGF-β +TGF-β-receptor-I (TGF-β-RI) kinase inhibitor SD-208 (right). Nuclei are stained with Hoechst 33258 (blue) and F-actin stained with Phalloidin (red). Below each image is a magnified image of the region indicated in red highlighting the localization of MGST1 in FB. Scale bar = 100 µm. **c.** Immunoblot showing MGST1 protein expression in control NF FB upon treatment with TGF-β and with co-application with SD-208. GAPDH was used as a loading control. **d.** Quantification of immunoblot data in b. One-way ANOVA (RM) was used with Tukey’s correction for multiple comparisons. *, p-value = 0.0193 & 0.0142; **, p-value = 0.0014. Error bars represent mean ± SD. ns, no significance.

Confocal imaging of immunostained cells showed that in control, consistent with its known localization to the endoplasmic reticulum (ER) and mitochondria, MGST1 was distributed in a granular pattern (Fig.4b) (27). Fibroblasts treated with TGF-β1 appeared larger with prominent F-actin stress fibres, consistent with myoFB differentiation. In a subset of cells, MGST1 had a fibre-like localization, possibly reflecting subcellular (e.g., mitochondrial) reorganization. In contrast, SD-208–treated cells adopted a more spindle-like morphology indicative of a less activated state, with F-actin arranged in more condensed patterns. Through Immunoblotting analysis, a 50% reduction in MGST1 was detected in FB treated for 4 days with TGF-β1 (Fig.4c and d). Co-treatment with SD-208 prevented this TGF-β-induced reduction in MGST1 and resulted in an increase in MGST1 above that in controls cells. This augmentation of MGST1 expression by SD-208 is explained by the reduction in baseline TGF-β receptor activation responsible for the low level of FB activation detected in FB cultures (28).

Together, these data indicate that loss of MGST1 is associated with FB activation and at least partly regulated by TGF-β signalling. Co-treatment with SD-208 attenuated the TGF-β1-induced reduction in MGST1 levels, further supporting the involvement of the TGF-β pathway in regulating MGST1 expression in activated FB.

### MGST1 knockdown FB exhibit some features of activation

We next examined whether the loss of MGST1 was sufficient for FB activation. To this end, we silenced MGST1 expression in human non-failing (NF) FB using siRNA (Fig.5a). In comparison to transfection with a non-targeting siRNA, transfection with an MGST1 targeting siRNA resulted in significant reduction in MGST1 expression at both protein and RNA levels (Fig.5b,c and Suppl.Fig.5a for full blot image). Notably, knockdown of MGST1 resulted in significant increase in mRNA expression levels of POSTN but not in COL1A1 and FAP (Fig.5d-f). While ACTA2 expression was not altered at mRNA level (Fig.5g), αSMA protein was significantly increased in MGST1 knockdown FB (Fig.5h and Suppl.Fig.5a for full blot image).

**Fig. 5.**
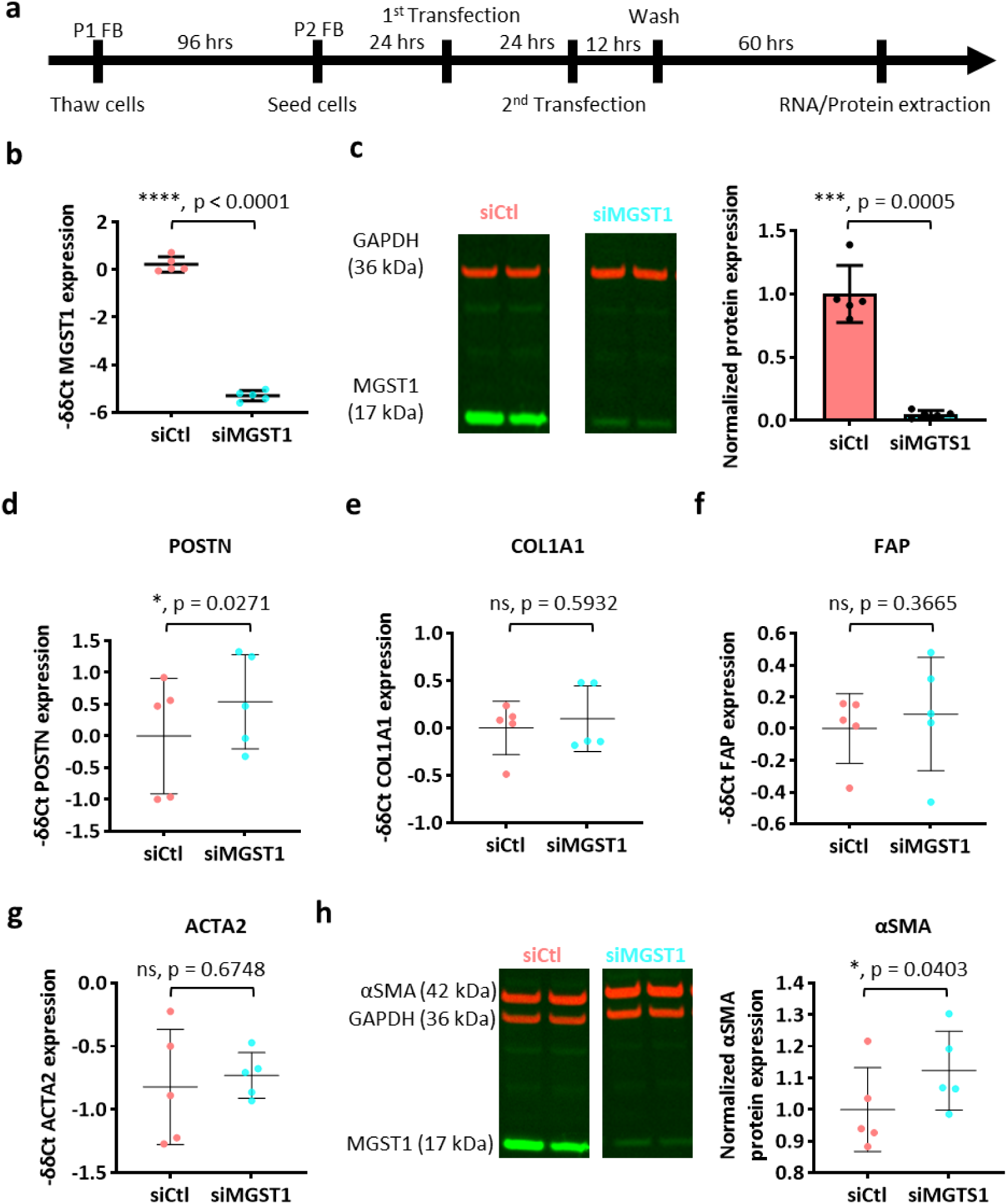
Loss of MGST1 is not sufficient for FB activation. **a.** Schematic representation of experimental protocol for MGST1 knockdown (KD). **b and c.** Relative mRNA (b) and protein (c) expression of MGST1 in control and MGST1 KD FB. Paired t-test. **d-g.** Relative mRNA expression level of fibrosis and FB activation related genes upon MGST1 KD vs control. Paired t-test. **h.** Relative protein expression level α-SMA upon MGST1 KD. Paired t-test. Error bars represent mean ± SD. ns, no significance.

We also investigated the effect of MGST1 knockdown on FB proliferation and viability. Using KI67 staining as a marker of proliferation (Suppl.Fig.5c), we did not detect any effect of MGST1 siRNA knockdown on the percentage of KI67 positive cells or the intensity of KI67 staining in the +ve nuclei (Suppl.Fig.5d & e). Additionally, there was no difference in the number of FB per field of view between the silenced and control group (Suppl.Fig.5f). MGST siRNA did not affect cell viability as determined by MTT assay (Suppl.Fig.5g).

Taken together, these data show that MGST1 silencing is not sufficient to induce full FB activation, but may contribute to the process once initiated as by TGF-β.

### MGST1 protects FB against oxidative stress

Given the proposed function of MGST1 to mitigate OS, we probed its role in in protecting FB against increased in ROS induced by cumene hydroperoxide (CHP; 25 µM), which increases cellular ROS through the generation of lipid radicals via cytochrome P450-catalyzed homolytic cleavage of the O–O bond, initiating lipid peroxidation and damaging cellular membranes (29) (Fig.6a). Live cell imaging using the ROS sensor CellROX Green was first performed to determine whether siRNA knockdown of MGST1 affected CHP induced increase in ROS in FB. MGST1 siRNA transfection had significantly higher CHP-induced ROS in comparison to FB transfected with a non-targeting siRNA control, indicating that the MGST1 acts to suppress cellular ROS levels (Fig.6b, c). The difference in CellROX fluorescence intensity between MGST1 silenced and control FB was apparent within ∼20 minutes from the addition of CHP, and remained for the 2 h duration of the experiment. Since an acute increase in ROS such as induced by CHP, can lead to cell detachment and death, we also measured FB cell death by counting the rounded FB after 2 h of CHP treatment. We observed significantly more rounded dead cells in the MGST1 silenced group compared to controls after 2 h of CHP treatment (Fig.6d).

**Fig. 6.**
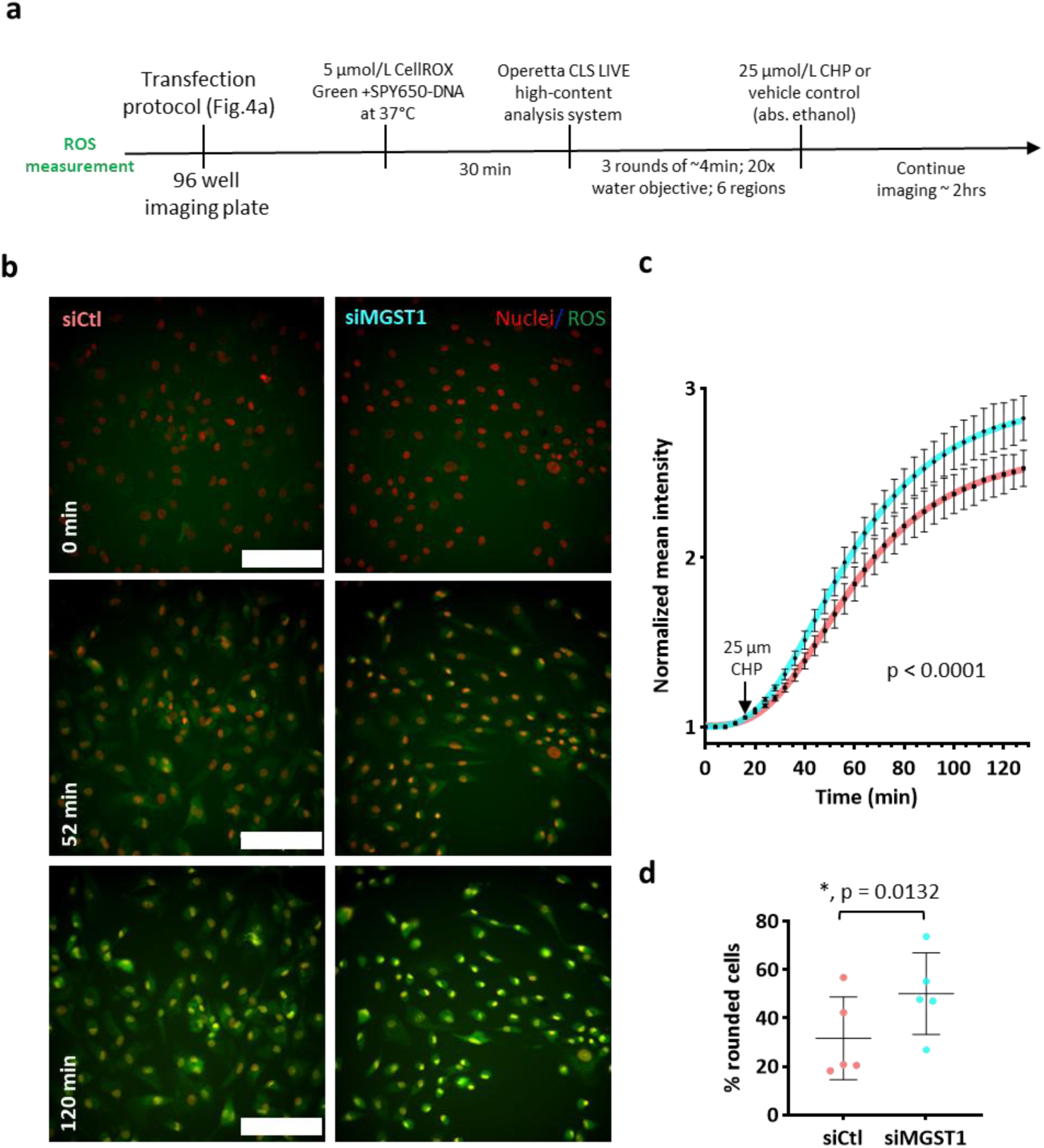
Loss of MGST1 enhances ROS production in response to stress. **a.** Schematic of time-line for cell staining and preparation for live cell imaging of ROS. **b.** Representative images of CELLRox fluorescence (green) live FB transfected with MGST1 (KD) or control siRNA and captured at the indicated timepoints. After 3 rounds of imaging (4 min interval each) to obtain a baseline, FB were treated with 25 µM CPH. Scale bar = 200 µm. **c.** Summary of data from b representing normalized mean intensity of CELLROX in all samples, conditions, and timepoints (4 min interval). Data was fit with non-linear dose-response curves. p-value corresponds to testing whether the 2 datasets can be fit by one curve or not. N=5. Error bars represent mean ± SEM. **d.** Quantification of rounded (detached/dead) cells after 2 h of CHP treatment. Paired t-test. Scale bar = 200 µm.

Since MGST1 performs its anti-oxidant activity in part via its utilization of reduced glutathione (GSH) to neutralize ROS (Fig.7a), we hypothesised that MGST1 knockdown would lead a reduction in GSH utilization and a consequent increase in its abundance after exposure to CHP. We therefore measured the effect of MGST knockdown on GSH levels. Single cell imaging of ThiolTracker (an indicator of reduced glutathione, GSH) detected no significant effect of MGST1 knockdown on basal levels of GSH (Fig.7b). GSH levels were however significantly higher in MGST1 silenced FB upon CHP treatment than in controls, indicative of heightened GSH availability, and supporting the modulation of peroxidase activity of MGST1 in FB (Fig.7c). Given that ROS contributes to cell death processes through peroxidation of lipids, and supported by the anti-oxidant activity of MGST1 identified above, we next probed whether MGST1 knockdown influenced lipid peroxidation in FB exposed to CHP. While no effect of MGST1 knockdown was detected at baseline (Fig.7d), after CHP treatment, oxidized lipid products were significantly greater in MGST1 knockdown FB than in controls (Fig.7e).

**Fig. 7.**
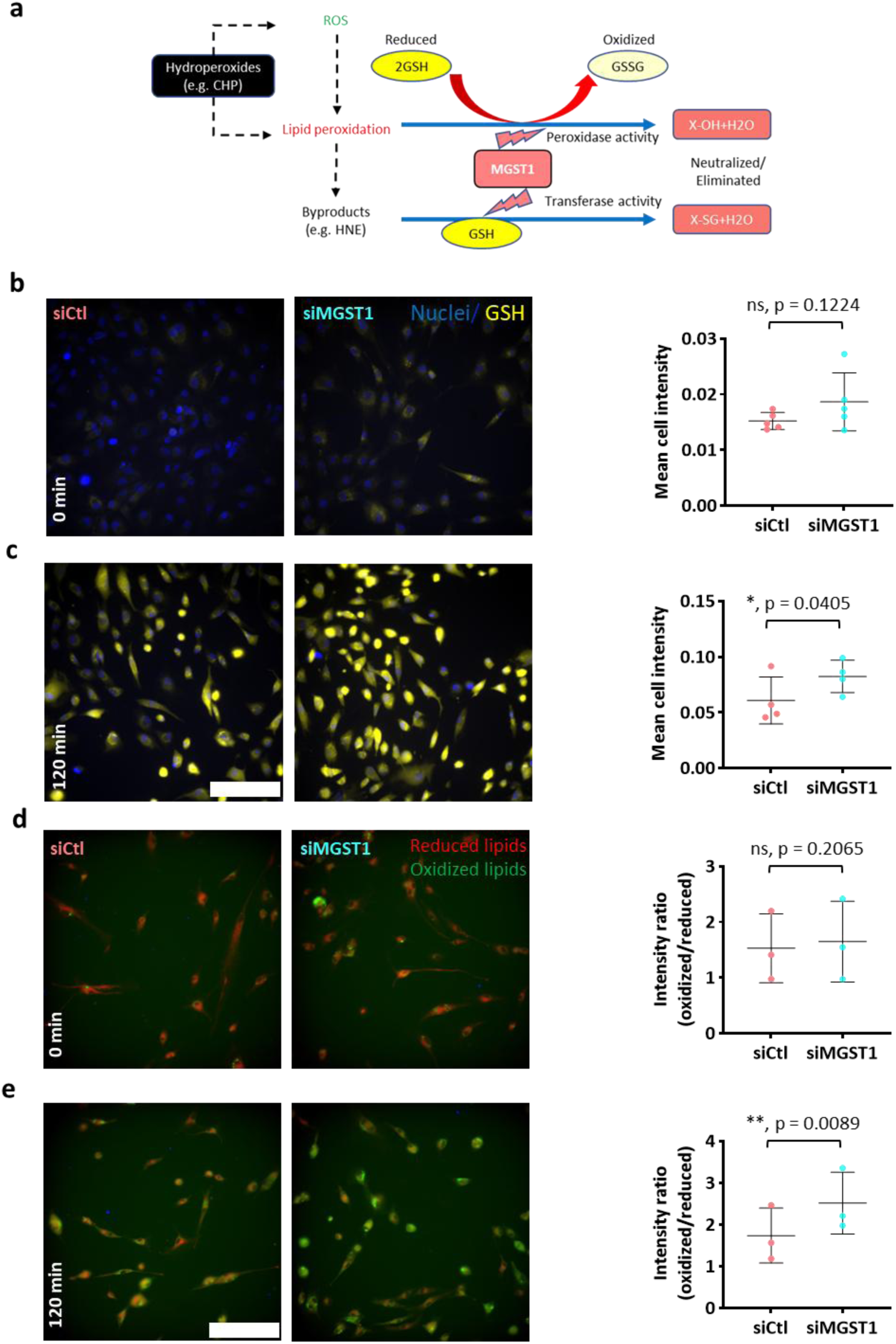
MGST1 knockdown FB exhibit reduced consumption of GSH and increased lipid peroxidation during oxidative stress. **a.** Schematic showing the possible actions of MGST1. GSH, reduced glutathione; GSSG, oxidized glutathione; HNE, Hydroxynonenal aldehyde, a byproduct of peroxidised lipids; CHP (cumene hydroperoxide). **b. Left,** Representative images of reduced Glutathione (GSH; yellow) in FB transfected with MGST1 siRNA or control at baseline before treatment with 25 µM CPH. Scale bar = 200 µm. **Right,** Quantification of normalized GSH intensity (showed in left) measured with ThiolTracker in MGST1 KD FB vs control. N=4 Paired t-test. **c. Left,** Representative images of reduced Glutathione (GSH; yellow) in MGST1 KD FB vs control after treatment with 25 µM CHP for 2 h. Scale bar = 200 µm. **Right,** Quantification of normalized GSH intensity (showed in left) measured with ThiolTracker in MGST1 KD FB vs control. N=4 Paired t-test. Error bars represent mean ± SD. ns, no significance. **d. Left,** Representative images of reduced and oxidized lipids (red/green) in MGST1 KD FB vs control at baseline before treatment with 25 µM CHP. Scale bar = 200 µm. **Right,** Quantification of reduced and oxidized lipids (red/green) ratio (showed in left) measured with Image-iT™ Lipid Peroxidation Kit in MGST1 KD FB vs control. N=3 Paired t-test. **e. Left,** Representative images of reduced and oxidized lipids (red/green) in MGST1 KD group vs control after treatment with 25 µM CHP for 2 h. Scale bar = 200 µm. **Right,** Quantification of reduced and oxidized lipids (red/green) ratio (showed in left) measured with Image-iT™ Lipid Peroxidation Kit in MGST1 KD group vs control. N=3 Paired t-test. Error bars represent mean ± SD. ns, no significance.

### HF FB are sensitised to death by Ferroptosis

MGST1 has previously been associated with ferroptosis (30,31), an iron-dependent regulated form of cell death driven by lipid peroxidation. Given this link and the observed protection against lipid peroxidation described above, we hypothesised that MGST1 was involved in the ferroptotic pathway in cardiac FB. To examine this hypothesis, we probed expression of ferroptosis related genes in our FB RNA-seq dataset. Kyoto Encyclopedia of Genes and Genomes (KEGG) analysis of the DEG identified in HF vs NF FB showed enrichment of the ferroptosis pathway in NF FB (Fig.8a). To further dissect this enriched ferroptosis geneset, we compared expression of genes that either drive or suppress ferroptosis, using a dedicated ferroptosis gene database (32). While ferroptosis-promoting genes remained largely unchanged between HF and NF FB, ferroptosis-suppressing genes were downregulated in HF (Fig.8b). We next examined how MGST1 related to these changes and found that MGST1 expression correlates with ferroptosis-inhibiting genes, but not with those that promote ferroptosis (Fig.8c). To provide additional evidence for a role for the ferroptosis pathway in FB in HF, we examined the expression of iron-handling genes, namely, the heavy and light chain subunits of ferritin (FTH1 and FTL respectively), which when complexed store excess cellular iron protecting cells from ferroptosis due to lower iron availability for ROS production. Both FTH1 and FTL were significantly reduced in HF compared to NF FB, suggesting a greater susceptibility to ferroptosis in the failing heart (Fig.8d).

**Fig. 8.**
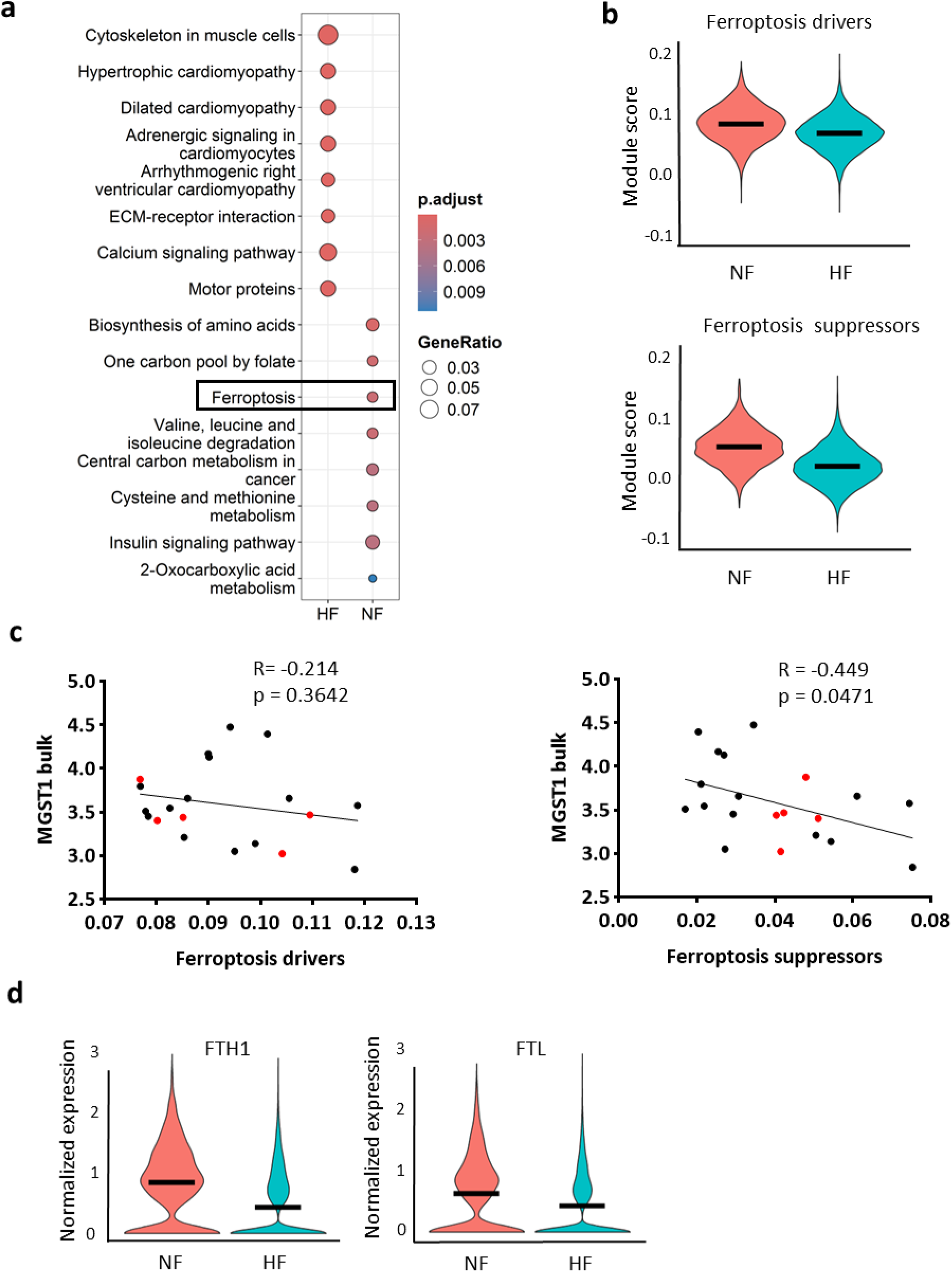
Ferroptosis suppressors are markedly downregulated in HF FB. **a.** The top 8 enriched KEGG pathways of HF FB DEGs compared to NF FB. **b.** Violin plots of module scores of ferroptosis driver (left) and suppressor (right) genes in HF and NF FB. **c.** Pearson correlation of MGST1 mRNA level with ferroptosis driver (left) and suppressor (right) genes module scores in each matched sample. Red dots represent samples from the ICM-scar region. **d.** Violin plots of normalized expression of FTH1 (left) and FTL (right) in HF and NF FB.

## Discussion

### Reduced MGST1 expression is linked to tissue fibrosis

We leveraged our recent snRNA-seq dataset from human failing (HF) and non-failing (NF) hearts to study alterations in the cellular ROS response genes, exploring a potential FB-specific pathway for tissue ROS modulation. We found that FB had the greatest reduction in expression of ROS-responsive genes in HF with FB-specific downregulation of MGST1. This finding aligns with the importance of ROS signalling in cardiac fibrosis (33), and resonates with reports linking fibrosis and MGST1 regulation in other cell types and organs, as well as in the heart.

MGST1 is strongly expressed in hepatocytes and several cancer types (27), and MGST1 has been shown to be decreased in liver cirrhosis (34), ovarian fibrosis (35,36), and keloid-associated skin fibrosis (37). Recent single-cell analyses identified MGST1 as a shared FB marker across multiple organs (heart, colon, bladder, skeletal muscle), suggesting a conserved role in FB stress responses and tissue remodelling (38). In the heart, a recent computational study using published human sequencing data identified a subset of FGF7⁺MGST1⁺ FB that are reduced in abundance post-MI (39). The study’s finding of MGST1 as a marker for a FB cluster in NF hearts is consistent with our observation of MGST1 as a marker for the FB2 cluster, which is abundant in NF and markedly decreases in HF (22).

MGST1 downregulation at both tissue mRNA and protein levels was also observed in diabetic cardiomyopathy together with ferroptosis (40), suggesting that its expression is governed by systemic factors during HF progression, such as inflammatory signalling, rather than disease-specific cues. Our intervention showing loss of MGST1 by TGF-β1 stimulation of healthy cardiac FB is consistent with inflammation as a general signal triggering its downregulation.

### FB specific decrease in anti-oxidant capacity exacerbates HF progression

Microsomal glutathione S-transferase 1 (MGST1) belongs to the MAPEG (membrane-associated proteins in eicosanoid and glutathione metabolism) family and consists of integral membrane proteins localized to the membranes of organelles (mainly the endoplasmic reticulum and mitochondria). The primary functions of MGST1 proteins include reducing lipid peroxidation and protecting intracellular membranes from oxidative stress at the expense of reduced glutathione (GSH) molecules by exhibiting both Glutathione S-transferase (GST) and glutathione peroxidase (GPx) activities (41,42). MGST1 also plays a protective role against ferroptosis by catalysing the conjugation of GSH to lipid peroxidation products to promote their degradation (30,31). We propose based on our data that loss of MGST1 contributes to HF progression by FB-specific loss of anti-oxidant capacity, with potentiation of fibrosis and of ferroptosis.

A first mechanism for potentiation of fibrosis is enhanced fibroblast activation. MGST1 is downregulated as part of the FB activation cascade, supported by the observed downregulation under TGB-β1 stimulation, but more importantly, low levels of MGST1 further enhance activation by modulating gene expression. Indeed, the increase of POSTN after selective knockdown of MGST1 fits into progression of NF FB substates into substates FB1 and FB4, the predominant clusters in HF (22). A second mechanism for potentiation is the decreased anti-oxidant capacity in FB, which would shift the balance towards higher ROS levels and a positive feedback on ROS-mediated FB activation.

In HF, increased ROS generation and decreased antioxidant capacity both contribute to higher levels of ROS (43). Measurements of ROS in tissue samples have identified the intracellular sources of ROS, with various contributions of mitochondria and NADPH oxidases depending on tissue and disease state (44,45). However, tissue measurements cannot discriminate which cells are mainly involved. Cardiomyocytes, given their mass, are likely a major source, but also endothelial cells, FB and inflammatory cells are involved in ROS balance at tissue level.

Our data complement current knowledge. Our observed downregulation of ROS-responsive genes suggesting that FB, though not the dominant ROS producers, serve as key antioxidant hubs. The anti-oxidant capacity of FB may not be limited to the FB themselves but may extend to protective effects on the cardiomyocytes, including protection against ferroptosis. The data implicate the ferroptosis GO pathway when comparing NF and HF FB, and show that ferroptosis suppressing genes (including MGST1) are downregulated in HF FB, rather than an upregulation in ferroptosis driver genes.

Mohr et al showed that FB protect cardiomyocytes through paracrine signalling and by mitigating the iron burden via intercellular interactions (15). The aforementioned study also showed that post-MI cardiomyocyte death is predominantly ferroptotic, and that resting and activated FBs (PDGFRα⁺, POSTN⁺) protect cardiomyocytes from ROS and ferroptosis via paracrine factors (e.g., IL-8, EGF) and direct iron handling through connexins (Cx43, Cx45), particularly in the border zone (15); FBs were also more ferroptosis-resistant, indicating superior resistance to oxidative stress.

### Translational perspectives

Clinically, trials of exogenous small-molecule antioxidants have largely underperformed (46,47). One explanation for this could be that intracellular antioxidant enzymes react orders of magnitude faster with ROS than small molecules and thus are much more effective in ROS scavenging (48). Accordingly, a therapeutic opportunity lies in augmenting endogenous antioxidant systems (enzymes and their substrates) rather than relying on extracellular scavengers or exogenous antioxidants.

Within this context, MGST1 is not the only antioxidant gene altered in HF, but its FB-enriched expression makes it a particularly promising target. To our knowledge, this is the first study to interrogate MGST1 expression, regulation, and function in cardiac FB, and our data indicate that MGST1 is downregulated in HF, aligning with independent high-throughput reports and reinforcing the rationale to restore endogenous antioxidant capacity in FB as a disease-modifying strategy.

## Conclusion

Our study highlights the role of MGST1 in regulating FB ROS and the impact of losing antioxidant capabilities in HF. Loss of MGST1 in HF is unique to FB, linked to FB activation. It impairs FB antioxidant capacity, exacerbates oxidative stress, and reduces resistance to ferroptosis. Loss of MGST1 also contributes to FB activation, altogether inducing a deleterious positive feedback on cardiac remodelling in human HF.

## Declarations

### Ethics approval

The study protocol conformed to the Helsinki declaration and was conducted according to national and European Union regulations on the use of human tissues and was approved by the ethical committee of UZ Leuven (S58824).

### Availability of data and materials

The datasets analysed during the current study are available in the Gene Expression Omnibus (GEO) repository, accession number GSE298023. All source data are provided within the paper and its supplementary information files. No custom code have been used in this study, however the code used for analysis are available from the corresponding author on reasonable request.

### Competing interests

The authors declare that they have no competing interests.

### Funding

This work was supported by project grants G0C7319N to KRS and FR, G0C6419N to HLR and G097021N to KRS and HLR from the FWO (Research Foundation Flanders) and by a C1 grant C14/21/093 to HLR, FR and KS from the research fund of KU Leuven. M.Y. received PhD stipend support from the Department of Cardiovascular Sciences, KU Leuven.

### Authors’ contributions

K.R.S., H.L.R., M.Y. designed experiments. M.Y., K.R.S., H.L.R. performed data curation and formal analysis. M.Y., O.R., A.P. performed experiments/acquired and/or analysed data. M.Y., K.R.S., H.L.R wrote the manuscript. F.R., M.Y., H.L.R. and K.R.S. contributed to Resources. All authors reviewed and edited the MS and approved the final submitted MS. K.R.S. and H.L.R. supervised the work.

## Acknowledgements

We thank Roxane Menten for technical assistance. We thank the UZ Leuven Transplant Team and Members of the UZ Leuven Cardiac surgery team for tissue procurement. We also thank the collaborators from Proteomics Leuven -Laboratory of Applied Mass Spectrometry (LAMaS) for their valuable contribution to the research presented in this paper.

## Non-standard Abbreviations and Acronyms

DCM: Dilated cardiomyopathy
DEG: Differentially expressed genes
FB: Fibroblasts
GO: Gene ontology
HF: Heart failure
ICM: Ischemic cardiomyopathy
myoFB: Myofibroblasts
MI: Myocardial infarction
NF: non-failing
OS: Oxidative stress
ROS: Reactive oxygen species
snRNA-Seq: single nucleus RNA sequencing
TGF-β (1): Transforming growth factor β (1)

## Notes

### Competing Interest Statement

The authors have declared no competing interest.

