## Supplementary Methods for "Loss of MGST1 during fibroblast differentiation enhances vulnerability to oxidative stress in human heart failure"

**snRNA-seq data analysis**

The single-nucleus RNA sequencing (snRNA-seq) data generation and preprocessing are detailed in Youness et al. ((1), in press). For the present study, only dilated cardiomyopathy (DCM) and ischemic cardiomyopathy (ICM) samples without apparent scar tissue were included. To assess oxidative stress activity, module scoring was performed in Seurat (v5.0.1) (2) using the *HALLMARK_REACTIVE_OXYGEN_SPECIES_PATHWAY* gene set obtained from MSigDB (https://www.gsea-msigdb.org/gsea/msigdb/human/geneset/HALLMARK_REACTIVE_OXYGEN_SPECIES_PATHWAY). Differential gene expression analysis was carried out at the pseudobulk level by summing raw counts per sample using AggregateExpression (Seurat) and testing with DESeq2 (v1.42.0) (3). DESeq2 was run with its default settings, including median-of-ratios normalization, negative binomial generalized linear modeling, and Wald tests for pairwise comparisons, with Benjamini–Hochberg correction for multiple testing. Pathway enrichment of differentially expressed genes was performed using KEGG implemented in clusterProfiler (v4.10.0) (4). Cell-type composition analysis was conducted at the per-sample level by quantifying relative cell proportions across groups. Pseudotime trajectory analysis of selected genes was performed using Monocle3 (v1.0.0) (5) as described in Youness et al (1). Bulk RNA-seq data of primary human cardiac fibroblasts treated with or without TGF-β1 were obtained from the publicly available dataset GSE225336 (6), and normalized expression of MGST1 was extracted for visualization. A curated list of ferroptosis-related genes was obtained from FerrDb V2 (7), a manually curated database of ferroptosis regulators and ferroptosis–disease associations.

**Fibroblast cell isolation and culture**

FB cell isolation and culture was done as described previously (8). Briefly, tissues samples were cut into small pieces using a surgical scalpel and washed with oxygen-rich, ice-cold Tyrode's solution (composition in mmol/L: NaCl 137, KCl 5.4, MgCl2 0.5, CaCl2 1.8, Na-HEPES 11.8, and glucose 10; pH 7.4). These tissues were enzymatically dissociated using collagenase-A (Roche, 10103586001, 0.229 U/mg) and protease XIV (P5147, SigmaAldrich, 0.35 mg/mL) in Normal Tyrode's solution in 2 rounds. The processed tissue was finely minced, and the resultant cell suspension was filtered (using 100 and/or 70 µm filters) to eliminate cellular debris. This suspension, rich in fibroblasts, was cultured in DMEM (Gibco, Thermo Fisher Scientific) enhanced with 10% fetal bovine serum (Gibco, Thermo Fisher Scientific) and 1% Antibiotic-Antimycotic (ThermoFisher Scientific). The culture underwent minimal passaging (p0-p2) to preserve the in vivo characteristics of the cells while still allowing for adequate growth for experimental studies. Purity of fibroblasts using this method was assessed previously in our group ((8), supplementary). In specific experiments, fibroblast activation was induced using TGF-β1 (10 ng/ml) with or without SD-208 (3 µmol/L; Sigma-Aldrich), a specific TGF-β-receptor-I (TGF-β-RI) kinase inhibitor. P2 FB were treated for 4 days (refresh media and treat again after 2 days). The number of hearts used for each experiment is summarized in the table below.

| Experiment | Tests | Number of hearts |
|---|---|---|
| TGF-β1/SD-208 treatment | qPCR-WB | N=4 |
| MGST1 KD experiment | qPCR-WB-ROS-Glutahtione | N=5 |
| MGST1 KD experiment | Lipid peroxidation | N=3 |

**Gene expression analysis**

Total RNA was isolated from tissue samples or cells using Qiazol (Qiagen) followed by phenol-chloroform extraction (according manufacturer's instructions). RNA was quantified using Nanodrop and cDNA was synthesized with Superscript III reverse transcriptase (ThermoFisher Scientific). The final cDNA reaction was diluted 1:20 in molecular biology grade water. RT-qPCR was performed in 6 µL reactions in 384-well format on a LightCycler® 480 Instrument II (Roche) using GoTaq RT-qPCR Mastermix (2X,Promega) with 1.5 µL cDNA and 300 nM concentration of each primer (Table 1). Gene expression was normalized to GAPDH. Primers were purchased from Integrated DNA Technologies.

*Table 1. Primers used for RT-qPCR*

| Gene | Forward primer | Reverse primer |
|---|---|---|
| MGST1 | TTGCCAATCCAGAAGACTGTGT | TGCGTACACGTTCTACTCTGTCA |
| COL1A2 | TGAGGGCAACAGCAGGTTCA | TCAGCACCACCGATGTCCAA |
| POSTN | GGAGACACACCCGTGAGGAA | CACTGAGAACGACCTTCCCTT |
| FAP | ATCCAGTGACCCACGCTCTG | CACCAATAAGGCAAGCACAGCA |
| ACTA2 | TGTGCTGGACTCTGGAGATG | GAAGGAATAGCCACGCTCAG |

**Immunofluorescence staining for fibroblast phenotyping in vitro and in situ and analysis**

Following the specified culture period, cells were fixed using 2% paraformaldehyde in PBS for 15 minutes at ambient temperature. Subsequently, the cells were permeabilized with 0.3% Triton X-100 for 30 minutes and blocked with 3% bovine serum albumin (BSA) for 60 minutes at room temperature. Cells were subsequently incubated overnight with primary antibodies diluted in blocking buffer. The antibodies and their dilutions used were as follows: Microsomal glutathione S-transferase 1 (MGST1) (1:100, ab131059, Abcam), KI67 (1:100, MAB7617-100, bio-techne). After washing three times for 5 minutes each, Secondary antibodies (1:500) were incubated for 1 hr at room temperature. After washing, F-actin stress fibres were stained using Rhodamine-Phalloidin (1:1000 dilution, ab176759, Abcam) coupled with AlexaFluor 647 conjugate. Nuclei were stained with Hoechst 33258 (1:1000). For tissue staining, membranes of tissue were stained with WGA647 (1:1000). Images were captured using Nikon A1R confocal microscope with 40X oil objective. Six to 9 images were taken per sample. For MGST1 quantification in cardiac tissue, exported TIFF images were analysed using CellProfiler (9) open-source software. To distinguish cardiomyocytes (CM) from non-cardiomyocytes (non-CM), cell membranes were first identified using WGA staining, and nuclei located within WGA-positive regions were detected. To define the non-CM compartment, a 10-pixel (4 µm) expansion was applied around the identified nuclei, thereby capturing the surrounding non-CM areas. Cardiomyocytes were excluded from analysis by restricting quantification to these expanded nuclear regions, ensuring that only non-CM signal was considered. MGST1 mean cell intensity per image was taken, and the average was used for each sample. A detailed workflow of this image analysis protocol is provided in Suppl.Fig.2d.

### Immunoblotting for protein expression analysis

Tissue lysates from left ventricular biopsies were prepared as previously described (8). Equal amounts of total protein homogenates were loaded onto 4–12% Bis-Tris gradient gels (ThermoFisher Scientific) for electrophoresis. Separated proteins were then transferred onto low fluorescence polyvinylidene difluoride (PVDF) membranes (Millipore). The membranes were blocked with Odyssey PBS blocking buffer (Li-COR Biosciences) and subsequently incubated overnight with primary antibodies diluted in 10% blocking buffer in TBS. The antibodies and their dilutions used were as follows: Microsomal glutathione S-transferase 1 (MGST1) (1:2500, ab131059, Abcam), α-SMA (1:5000, A2547, Sigma-Aldrich). After washing off excess antibodies with TBST for 1 hour, the membranes were incubated with the corresponding secondary antibodies (Li-COR 800CW or Alexa Fluor 680, ThermoFisher Scientific). Immunofluorescence detection was performed using an infrared imaging system (Odyssey CLx, LI-COR). Band intensities were quantified using Image Studio software, with the intensities of the proteins of interest normalized to GAPDH (1:5000, G8795, Sigma-Aldrich) in the same lane, which served as a reference standard. For protein analysis in isolated cells, the cells were lysed in RIPA buffer, and whole cell lysates were quantified using a BCA assay. Subsequent protocols followed the procedures described above.

### Sample Preparation for Mass Spectrometry Analyses

Cell pellets were lysed in RIPA buffer. A total of 10 µg of protein was mixed with 10% SDS and 50 mM triethylammonium bicarbonate (TEAB), and protein digestion was performed using the S-Trap kit (Protifi) according to the manufacturer’s instructions. Following cell lysis and buffer addition, samples were reduced with 5 mM tris(2-carboxyethyl)phosphine (TCEP), alkylated with 20 mM iodoacetamide (IAA), and denatured using 2.5% (v/v) phosphoric acid.

Subsequently, samples were diluted with 165 µl of binding buffer (90% methanol in 100 mM TEAB) and loaded onto S-Trap micro columns. Columns were centrifuged for 30 seconds at 10,000 × g, followed by three washes with 150 µl of binding buffer. Trypsin was added at a 1:10 enzyme-to-protein ratio in 20 µl of 50 mM TEAB, and digestion was carried out overnight at 37°C. Peptides were sequentially eluted from the column using 40 µl of elution buffer 1, 2 and 3 and (50 mM TEAB, 0.2% formic acid (FA) in water, and 40 µl water/acetonitrile (ACN) (50/50, v/v), respectively. Eluted peptides were dried completely using vacuum centrifugation prior to LC-MS/MS analysis.

**LC-MS/MS: Sample Acquisition via DIA and Data Analysis**

Tryptic peptides were reconstituted in 0.5% acetonitrile (ACN) and 0.1% formic acid (FA), and 300 ng of peptide material was loaded onto Evotips according to the manufacturer's guidelines (Evosep). Peptide separation and acquisition were performed using an Evosep One liquid chromatography system (Evosep) coupled to a ZenoTOF 7600 mass spectrometer (SCIEX), operating under Sciex OS software (version 3.3 or later). Peptides were separated using the 30 samples per day (30 SPD) method, which employs a 44-minute gradient and the EV1109 Performance column (ReproSil Saphir C18, 1.5 µm particle size, 8 cm × 150 µm; Dr. Maisch) maintained at 40 °C. The system was configured with the low micro electrode and operated at flow rates of 1–10 µL/min. Mobile phases consisted of 0.1% FA in LC–MS-grade water (buffer A) and 0.1% FA in ACN (buffer B). Mass spectrometry acquisition was conducted in SWATH data-independent acquisition (DIA) mode using the ZenoTOF with an OptiFlow

Turbo V ion source. Source parameters were as follows: spray voltage of 4500 V, gas 1 at 20 psi, gas 2 at 10 psi, curtain gas at 40 psi, and source temperature set to 100 °C. The Zeno SWATH method used 65 variable-width isolation windows covering the 400–1250 m/z mass range. TOF-MS scans ranged from 400–1250 m/z with a 25 ms accumulation time, while TOF-MS/MS scans ranged from 230–1400 m/z with a 20 ms accumulation time. Dynamic collision energy was applied with a charge state of 2, and Zeno pulsing was enabled. Raw data files (.wiff format) were processed using Spectronaut (version 19) in directDIA+ (deep) mode. Spectra were searched against the Human reference proteome (UniProt ID: UP000005640; https://www.uniprot.org/proteomes/UP000005640 ) using default Spectronaut parameters: precursor and protein Q-value thresholds set to 0.01; MS2 area used for quantification; normalization strategy set to "Automatic"; and default mass tolerance settings. Variable modifications included oxidation of methionine and N-terminal acetylation, while carbamidomethylation of cysteine was set as a fixed modification. Protein quantification reports were exported and subjected to downstream analysis.

**MGST1 Gene Knockdown**

Human cardiac fibroblasts isolated from non-failing hearts were cultured in Dulbecco's Modified Eagle Medium (DMEM) supplemented with 10% fetal bovine serum (FBS) and 1% Antibiotic-Antimycotic. For MGST1 gene knockdown, fibroblasts were transfected with siRNA targeting MGST1 (4390824, s8757, Fischer Scientific) using Lipofectamine™ RNAiMAX Transfection Reagent (ThermoFisher Scientific) according to the manufacturer's protocol. Briefly, cells were seeded in respective plates at a density of 40k cells per $cm^2$ and incubated overnight. The following day, siRNA-Lipofectamine™ RNAiMAX complexes were prepared by diluting siRNA targeting MGST1 (**s8757**, ThermoFisher Scientific) or siRNA negative control (4390843, ThermoFisher Scientific) and Lipofectamine™ RNAiMAX separately in Opti-MEM™ Reduced Serum Medium (ThermoFisher Scientific), then combining them and incubating for 5 minutes at room temperature. The complexes were added to the cells to achieve a final siRNA concentration of 20 nM. After 24 hours, the medium was replaced with fresh DMEM containing 10% FBS and new complexes were added for another 12 hrs, which were then replaced by normal growth media. Cells were incubated for 60 hours post-transfection before further analysis.

**Oxidative stress measurement and analysis in cardiac tissue and primary human fibroblasts**

To measure oxidative in cardiac tissue, cleaned cardiac tissue was cut into small pieces using surgical scissors and embedded in OCT Compound, then immediately placed in methylbutane container within larger LN2 container for snap freezing. Frozen sections, 10 µm thick, were cut using a cryostat at −20 °C and mounted onto histological slides, which were stored at −80 °C. A fresh 5 µM DHE (D11347, ThermoFisher Scientific) staining solution was prepared by dissolving 1 mg DHE in 317 µL dimethyl sulfoxide (DMSO) to create a 10 mM stock solution, which was then diluting 75 µL with 150 mL Milli-Q pure H2O. Hoechst 33258 was added (1:1000) to stain nuclei. Slides were rinsed in pure H2O for 30 seconds to remove OCT Compound, then immediately placed in the DHE staining solution and incubated for 20 minutes at room temperature, protected from light. After staining, slides were washed twice in deionized H2O for 1 minute each and kept in deionized H2O. Antifade mounting medium were added to the slides and a coverslip was placed and secured with nail polish at the corners.

Fluorescence imaging was performed immediately using a fluorescence microscope with red and blue excitation filters to visualize DHE-derived 2-OH-E+ and nuclei, respectively. Images were captured using Nikon A1R confocal microscope with 40X oil objective, with exposure times optimized to minimize background. Fluorescence images of cardiac tissue sections were acquired using identical laser power, exposure time, sensitivity, and resolution, selecting at least 5 different fields for each sample.

TIFF images were analysed for mean nuclei intensity using CellProfiler open-source software. Nuclei were identified with the Hoechst 33258 blue channel, and DHE was quantified using red channel. As for quantifying percent DHE staining per image, Fiji image J open-source software was used via a macro for batch processing. Thresholding was performed (min = 500) and area of DHE was recorded.

For measuring reactive oxygen species (ROS) in cells, CellROX Green reagent (C10444; Invitrogen), a fluorogenic probe, was used according to the manufacturer's instructions with some modifications. Briefly, after MGST1 silencing, cells cultured in 96-well plate were incubated with final concentration 5 µmol/L CellROX Green for 30 minutes at 37°C. SPY650-DNA (1x, SpiroChrome) was used to stain live nuclei. Cells were then imaged before and after the treatment of cumene hydroperoxide (CHP, 25 µmol/L) or vehicle control (absolute ethanol). For image acquisition, an Operetta CLS LIVE high-content analysis system was used equipped with a large format sCMOS camera in combination with a 20x water objective. The setup was controlled by Harmony software (Perkin Elmer Inc. USA). Six images from different (consistent) areas in each well was taken.

Single channel images were exported as TIFF files, and post processing/imaging analysis was done using CellProfiler open-source software. Since CellROX Green binds DNA upon oxidation, we followed ROS by measuring the oxidized DNA in the nuclei of each Fibroblast cell. Nuclei were identified with the SPY650-DNA 647 channel, and ROS was quantified using CellROX Green channel. Mean cell intensity parameter was used to analyse the data. After baseline correction, fluorescence was displayed as f/f0.

**Glutathione detection**

Cellular GSH levels were monitored and analysed using the ThiolTracker Violet GSH detection reagent (T10096; Invitrogen) according to the manufacturer's protocol. Briefly, after MGST1 silencing and treatment with CHP or vehicle (CHP, 25 µmol/L) for 2 hours, cells were rinsed with D-PBS were incubated with final concentration 20 µmol/L of ThiolTracker for 30 minutes at 37°C. Then the D-PBS was changed with DMEM media with no phenol. For image acquisition, an Operetta CLS LIVE high-content analysis system was used equipped with a large format sCMOS camera in combination with a 20x water objective. The setup was controlled by Harmony software (Perkin Elmer Inc. USA). Fourteen images from different (consistent) areas in each well was taken. Single channel images were exported as TIFF files, and post processing/imaging analysis was done using CellProfiler open-source software. Fibroblast cells were identified using GSH channel and mean cell fluorescence intensity were measured. Median of these cells' intensity were visualized.

**Lipid peroxidation detection**

Lipid peroxidation was monitored using the Image-iT® Lipid Peroxidation Kit (C10445; Invitrogen) according to the manufacturer's protocol. Briefly, after MGST1 silencing and treatment with CHP (25 µmol/L) or vehicle control (absolute ethanol) for 2 hours, cells were incubated with 10 µM Image-iT® Lipid Peroxidation Sensor for 30 minutes at 37 °C, rinsed with PBS, and imaged immediately. An Operetta CLS LIVE high-content analysis system (Perkin Elmer Inc., USA) equipped with a large format sCMOS camera and 20x water objective was used for acquisition, capturing fourteen images from consistent areas per well. Upon oxidation, the reagent shifts fluorescence emission peak from ~590 nm (red) to ~510 nm (green). Single channel images were exported as TIFF files, and post-processing/quantification of the red/green ratio was performed in CellProfiler. Median per-cell ratios were visualized to quantify lipid peroxidation levels.

**Statistics and reproducibility**

GraphPad Prism software (10.2.3) was used for statistical testing. The test used for each experiment is mentioned in figure legends. P-value < 0.05 was considered statistically significant, and each p-value and test used were reported in the figures/ figures legend. Analysis of sequencing data was performed in R (4.3.2). No customized code was used in this analysis.
