## Supplementary material for "Loss of MGST1 during fibroblast differentiation enhances vulnerability to oxidative stress in human heart failure": Suppl.Fig.

**Supplementary Figures**

**Supplementary Fig.S1**

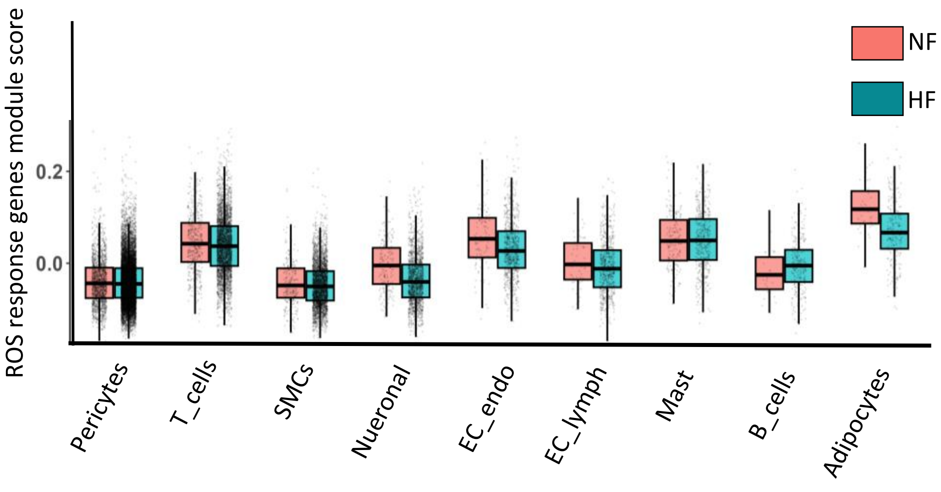


**Supplementary Fig.S1**
Scoring of ROS response gene expression in low abundant cardiac cell types not described in Fig 1a. Genes taken from the HALLMARK_REACTIVE_OXYGEN_SPECIES_PATHWAY (M5938).

### Supplementary Fig.S2

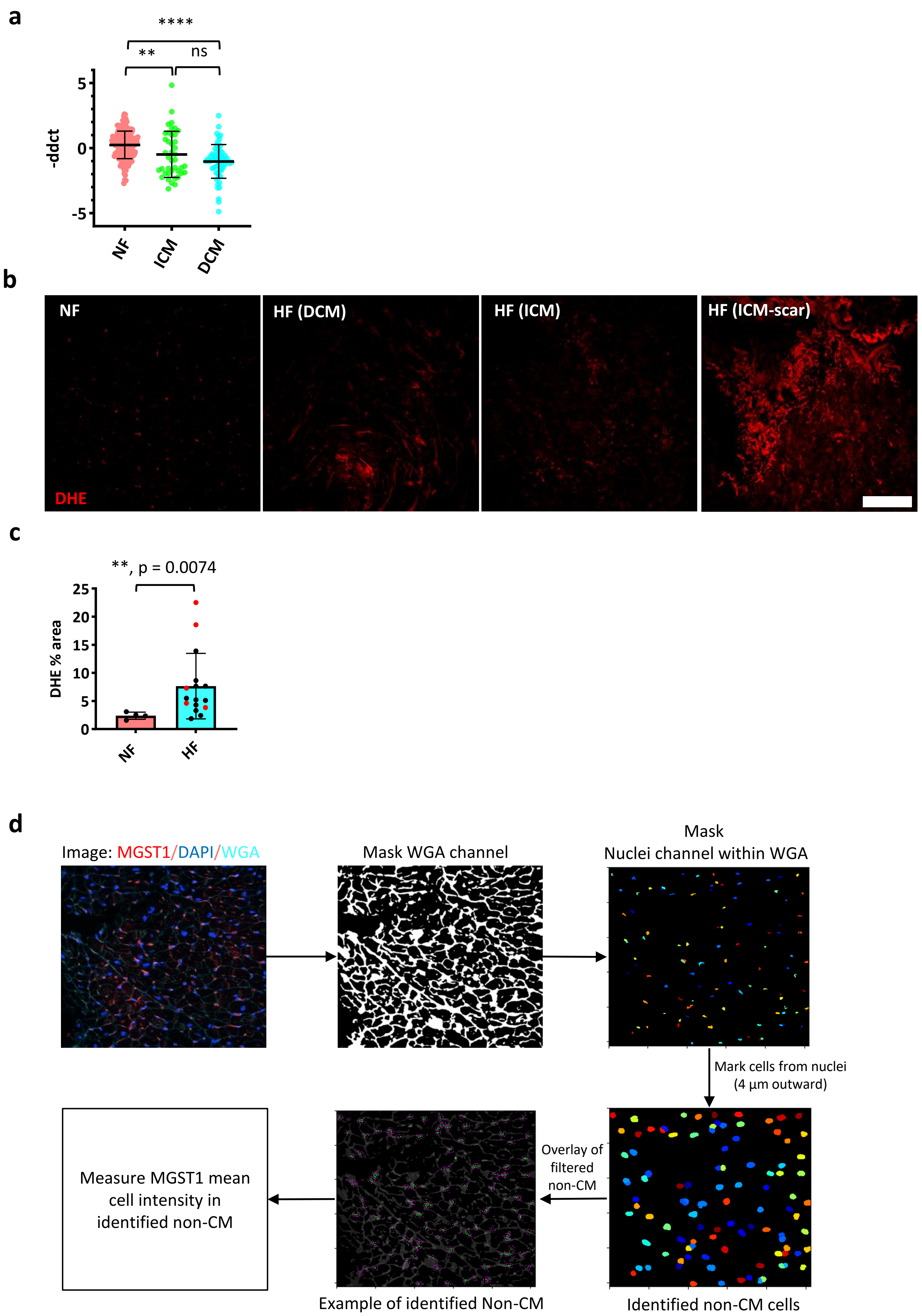

**Supplementary Fig.S2**

**a**. Relative mRNA expression of MGST1 in human cardiac tissue demonstrating decrease of MGST1 expression in HF (ICM and DCM) compared to NF. One-way ANOVA. p-value: **, 0.0028 ,*** < 0.0001. p-value (ICM vs DCM) = 0.0810. Multiple comparison was corrected by Tukey's test.

**b**. Detection of superoxide in myocardium by DHE staining. Scale bar = 100 µm.

**c**. Qauntification of DHE percentage of area covered in NF and HF myocardium. Mann-Whitney test. Error bars represents mean ± SD.

**d**. Schematic showing the imaging analysis pipeline for MGST1 quantification in tissue.

#### Supplementary Fig.S3

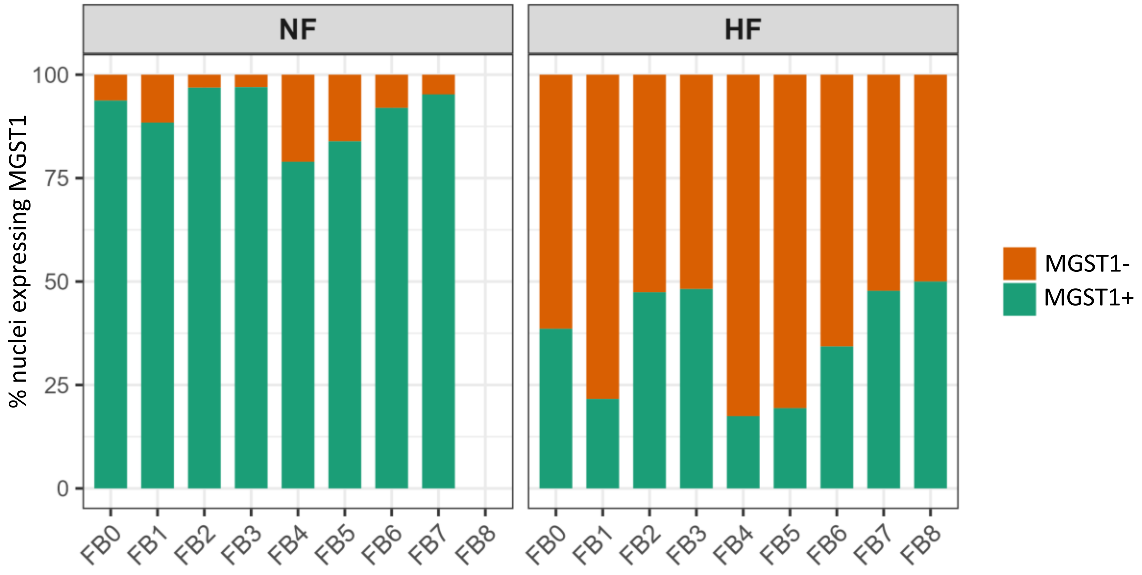


**Supplementary Fig.S3**
Stacked Barplot showing the percentage of nuclei expressing MGST1 in NF and HF FB states.

#### Supplementary Fig.S4

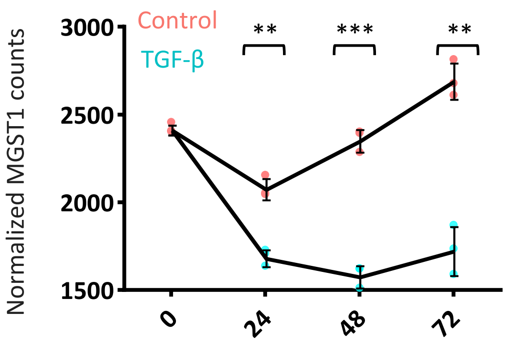


**Supplementary Fig.S4**
Normalized MGST1 counts of MGST1 in human primary Fb during different time points and upon TGFb treatment. Data was extracted and analyzed from RNAseq dataset of Nauffal et al (GSE225336). 2-way ANOVA test was used with Šidák correction for multiple comparisons. p-value (24 hr) = 0.0045, p-value (48 hr) = 0.0005, p-value (72 hr) = 0.0038.

**Suppl.Fig.5 MGST1 knock-down in NF FB does not lead to FB proliferation**

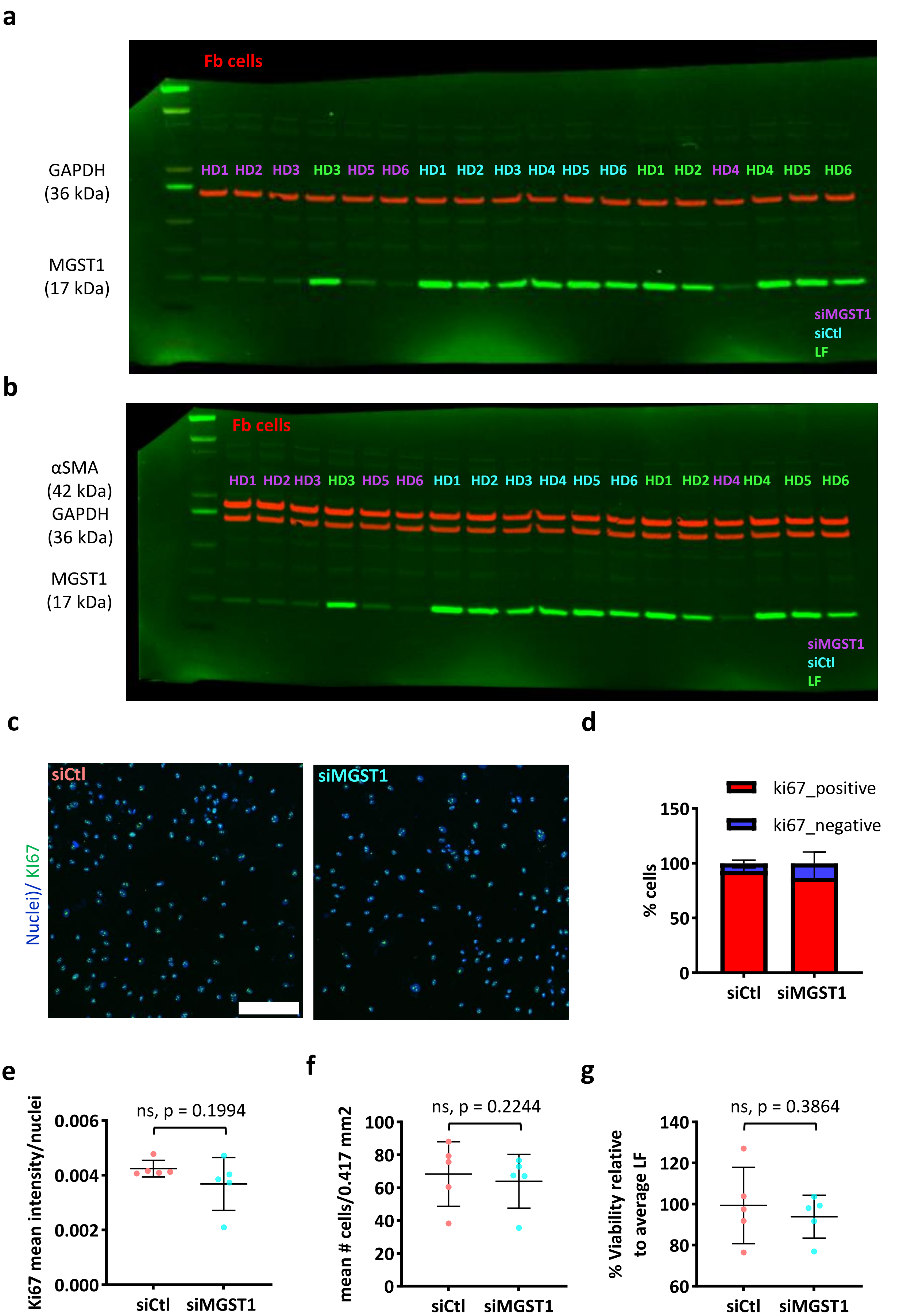

**Fig.5 MGST1 KD affects Fb differentiation but not Proliferation**

**a**. Full immunoblot image of representative data shown in Fig.5c.

**b**. Full immunoblot image of representative data shown in Fig.5h. Note: same blot used in (a) but re-probed with α-SMA antibody.

**c.** KI67 fluorescent staining (green) in MGST1 KD group vs control. Nuclei stained with Hoechst 33258 (blue). Scale bar = 200 µm.

**d and e.** Quantification of KI67 staining using % cells positive of KI67 (d) and mean nuclei intensity (e) in MGST1 KD group vs control. Paired t-test. p-value = 0.2432 for (d).

**f.** Number of Fb cells per 1 mm2 of field showing nuclei number in MGST1 KD group vs control. Paired t-test.

**g.** Percentage viability in MGST1 KD FB relative to control as measured by MTT assay. Paired t-test. Error bars represents mean ± SD. ns, no significance.
